# Not all branches are equal: Branch Length Evaluation for Phylogenetic Diversity analysis

**DOI:** 10.1101/2025.01.11.632535

**Authors:** Daniel Rafael Miranda-Esquivel

**Affiliations:** Escuela de Biología. Universidad Industrial de Santander, Bucaramanga, Colombia

**Keywords:** PD, null model, branch length, sensitivity analysis

## Abstract

Phylogenetic Diversity (PD) is a crucial metric in evolutionary and conservation studies; however, its dependence on branch-length information can be a source of uncertainty. Branch Length Evaluation for Phylogenetic Diversity analysis (BLEPD) is a free and open-source R program used to quantify the uncertainty in PD calculations caused by varying branch length information. It offers two main approaches: resampling branch lengths within the phylogenetic tree, and modifying individual branch lengths. By comparing the initial and selected areas after resampling or modification, BLEPD allows researchers to assess the sensitivity of PD to these variations. This analysis of branch length sensitivity provides valuable insights into the robustness of PD results and their dependence on this information. BLEPD may improve the interpretation of PD analyses in evolutionary, conservation, and ecological studies.

## Introduction

Vane-Wright et al. (1991) proposed a phylogenetic approach to conservation, emphasizing the importance of preserving evolutionary history, because prioritizing the conservation of phylogenetically distinct species can help maximize the protection of evolutionary history and functional diversity.

Phylogenetic diversity (PD) is a measure of biodiversity that considers the evolutionary relationships among species within a given set, such as a community or geographic region (Faith, 1992). Unlike species richness, which simply counts the number of species, PD accounts for evolutionary history. It is calculated as the sum of the branch lengths in the phylogenetic tree that connects the species within the set (Faith, 1992). By incorporating evolutionary history, PD provides a more comprehensive assessment of biodiversity, recognizing that species with distant evolutionary relationships contribute more to overall diversity than closely related species. PD can be used as a proxy for functional diversity, because closely related species are more likely to share similar traits. Therefore, a higher PD should correspond to a greater functional diversity (Davies and Buckley, 2011).

PD is also considered to be a measure of evolutionary potential, which is the capacity of a group of species to adapt to environmental changes (Winter et al., 2013). The loss of phylogenetically distinct species could mean losing unique adaptations and reducing the ability of an ecosystem to respond to future challenges.

Various methods exist for calculating and estimating PD. Traditional methods rely on the construction of phylogenetic trees, whereas alternative methods use statistical approaches to estimate the PD from incomplete or uncertain data (Cardoso et al., 2014).

A traditional approach to evaluating the structure of PD involves using null models to assess the significance of observed values. These null models typically randomize aspects of the data, such as species labels, richness, or occurrence patterns, while maintaining certain ecological properties [see functions **ses.pd** and **randomizeMatrix** within the picante package (Kembel et al., 2010)]. However, these traditional null models primarily focus on community-level properties and do not capture the impact of the topology or branch lengths on the observed PD values.

### BLEPD: Branch Length Evaluation for Phylogenetic Diversity analyses

BLEPD, a free and open-source R program available on GitHub (github.com/Dmirandae/blepd), evaluates branch length uncertainty in Phylogenetic Diversity (PD) analysis. It offers two independent approaches: resampling branch lengths and modifying individual branches. This allows researchers to identify influential branches and assess the distribution of PD across the phylogeny. By analyzing how PD values change with these modifications, BLEPD provides insights into the sensitivity and confidence of PD estimates, enabling a deeper understanding of the factors shaping phylogenetic diversity, including the influence of individual branches, clades, and the evenness of PD distribution. This leads to deeper insights into the relationship between evolutionary history and biodiversity patterns.

For consistency, functions are in **bold** typeface, arguments are in *italics*, and the values for the argument are underscored.

Resampling Branch Lengths: **swapBL**

The **swapBL** function within the BLEPD package is based on a null model approach designed to assess the sensitivity of PD calculations to uncertainties in branch-length estimates. The function employs three different branch length resampling models (argument *model*) within a specified scope of branches (*branch*: <u>terminals</u>, <u>internals</u>, or <u>all</u>):

simpleswap: Swaps the lengths of two randomly selected branches.

allswap: Random permutation of all branch lengths.

uniform: Replaces the branch lengths of the chosen nodes with random values drawn from a uniform distribution bounded by the minimum and maximum values of the original branch lengths.

For each resampling, the function iterates a specified number of times. In each iteration, the tree is modified according to the selected model. Subsequently, the PD value for the modified tree is calculated and the set with the highest PD is recorded. After all iterations are completed, the function summarizes the results by creating a data frame that records the frequency of each set chosen as having the highest PD across all iterations.

The comparison of these results with the set(s) selected in the initial tree is key to evaluating the evenness of the branch length distribution and how variations in that distribution affect PD calculations, thus informing the confidence in the identified sets of higher diversity. A general guideline is that a smaller difference between the observed selected set(s) and the randomized selected set indicates greater confidence in the initial selection. A difference of more than 5% can be considered a preliminary threshold for identifying potentially low confidence in the results.

Modifying Individual Branch Lengths: **evalBranch**

The **evalBranch** function was designed to evaluate the effect of changing a specific branch length as a measure of the stability of the results. First, the function calculates the initial PD value using the provided tree and the distribution data. If the *approach* is set to <u>lower</u>, the branch length of the target branch is set to zero. Conversely, if the *approach* is <u>upper</u>, the maximum permissible branch length is calculated based on a multiplier (the default value is 1.01) and the total tree length. The tree is subsequently modified with this new branch length and the function recalculates the PD value using the modified tree. The sets that exhibit the highest PD values in both the initial and modified scenarios are compared. If the sets with the highest PD values differ, the algorithm enters an iterative refinement approach that utilizes a bisection search to find either the minimum (“lower”) or the maximum branch length (“upper”) that causes a change in the selected area.

This analysis provides useful insights into the sensitivity of PD to changes in individual branch length. By comparing the “lost lengths” (for decreasing branch lengths) or “gained lengths” (for increasing branch lengths) across different branches, we can assess the overall sensitivity of PD results. Although there is no established threshold for identifying potentially unstable results, branches requiring larger “lost” or “gained” lengths to cause a shift in the optimal set indicate greater PD stability.

## Examples

### A Four Taxon Example

Here, I explore the potential influence of a single long branch on PD analysis in a simple four-taxon symmetrical topology with terminals equally and balancedly distributed across sets (Figure 1) with two scenarios: equal and unequal branch lengths.

**Figure 1.**
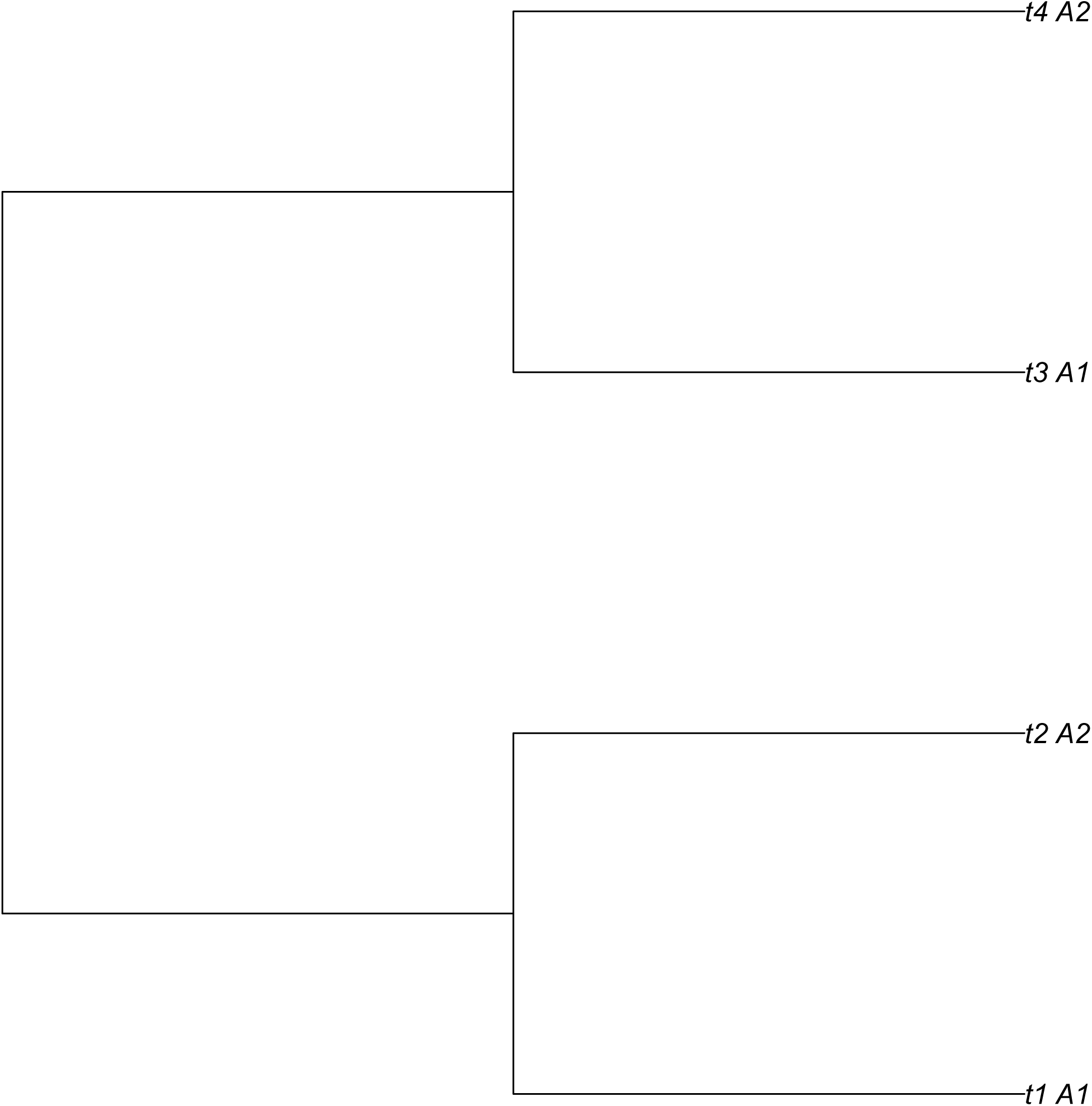
Four terminals (t1-t4) and their distribution into two areas (A1 and A2).

When all branch lengths are equal, the initial PD outcome is a tie for both sets due to the symmetry in path lengths. Consequently, any branch swap, regardless of the model used, has no effect on PD, and changes in internal branch lengths do not affect PD values. However, even slight variations in terminal branch lengths can impact PD outcomes, making the final result highly sensitive. Despite the even distribution of branch lengths, this arrangement is unstable as small changes in length can lead to substantial PD variations and ultimately influence the selection of the optimal set (Tables 1 and 2).

**Table 1.** Results for SwapBL 4-Terminal, 2-Area Example. The data shows the impact of swapping branch lengths on the selected area with the highest PD as the percentage of times each area or range (A1, A2 or A1A2) was selected as optimal under different scenarios. Conventions: ‘terminals’: Swapping only terminal branches. ‘internals’: Swapping only internal branches. ‘all’: Swapping both terminal and internal branches. ‘allswap’: Swapping all branch lengths randomly. ‘uniform’: Resampling branch lengths from a uniform distribution. l_t1t2: Internal branch length. l_t1: Terminal branch length. PD-A1: Phylogenetic Diversity of Area 1. PD-A2: Phylogenetic Diversity of Area 2.

| branchSwapped | model | $l_{t1t2}$ | $l_{t1}$ | A1 | A1A2 | A2 | PD-A1 | PD-A2 |
| --- | --- | --- | --- | --- | --- | --- | --- | --- |
| terminals | allswap | 1 | 1 | 0 | 100 | 0 | 4 | 4 |
| internals | allswap | 1 | 1 | 0 | 100 | 0 | 4 | 4 |
| all | allswap | 1 | 1 | 0 | 100 | 0 | 4 | 4 |
| terminals | uniform | 1 | 1 | 0 | 100 | 0 | 4 | 4 |
| internals | uniform | 1 | 1 | 0 | 100 | 0 | 4 | 4 |
| all | uniform | 1 | 1 | 0 | 100 | 0 | 4 | 4 |
| terminals | allswap | 2 | 1 | 0 | 100 | 0 | 5 | 5 |
| internals | allswap | 2 | 1 | 0 | 100 | 0 | 5 | 5 |
| all | allswap | 2 | 1 | **33.28** | 33.54 | 33.18 | 5 | 5 |
| terminals | uniform | 2 | 1 | 0 | 100 | 0 | 5 | 5 |
| internals | uniform | 2 | 1 | 0 | 100 | 0 | 5 | 5 |
| all | uniform | 2 | 1 | **50.66** | 0 | 49.34 | 5 | 5 |
| terminals | allswap | 1 | 2 | **49.77** | 0 | 50.23 | 5 | 4 |
| internals | allswap | 1 | 2 | 0 | 100 | 0 | 5 | 4 |
| all | allswap | 1 | 2 | **32.76** | 33.29 | 33.95 | 5 | 4 |
| terminals | uniform | 1 | 2 | **50.56** | 0 | 49.44 | 5 | 4 |
| internals | uniform | 1 | 2 | 0 | 100 | 0 | 5 | 4 |
| all | uniform | 1 | 2 | **50.24** | 0 | 49.76 | 5 | 4 |

**Table 2.** Results of the evalBranch function for the 4-Terminal, 2-Area Example. This table shows the impact of modifying individual branch lengths on the selected area with the highest Phylogenetic Diversity (PD). node: The node identity of the branch being modified. initialArea: The area with the highest PD in the initial tree. FinalArea: The area with the highest PD after the branch length modification. Approach: The type of branch length modification (“lower” for decreasing to zero, “upper” for increasing to the maximum). %Delta: The percentage change in branch length. l_t1t2: The length of the internal branch connecting terminals 1 and 2. l_t1: The length of the terminal branch leading to terminal 1. PD-A1: Phylogenetic Diversity of Area 1. PD-A2: Phylogenetic Diversity of Area 2.

| node | initialArea | FinalArea | Approach | %Delta | l_t1t2 | l_t1 | PD-A1 | PD-A2 |
| --- | --- | --- | --- | --- | --- | --- | --- | --- |
| [t1] | A1A2 | A2 | lower | -1 | 1 | 1 | 4 | 4 |
| [t2] | A1A2 | A1 | lower | -1 | 1 | 1 | 4 | 4 |
| [t3] | A1A2 | A2 | lower | -1 | 1 | 1 | 4 | 4 |
| [t4] | A1A2 | A1 | lower | -1 | 1 | 1 | 4 | 4 |
| [t1] | A1A2 | A1 | upper | 1 | 1 | 1 | 4 | 4 |
| [t2] | A1A2 | A2 | upper | 1 | 1 | 1 | 4 | 4 |
| [t3] | A1A2 | A1 | upper | 1 | 1 | 1 | 4 | 4 |
| [t4] | A1A2 | A2 | upper | 1 | 1 | 1 | 4 | 4 |
| [t1] | A1A2 | A2 | lower | -1 | 2 | 1 | 5 | 5 |
| [t2] | A1A2 | A1 | lower | -1 | 2 | 1 | 5 | 5 |
| [t3] | A1A2 | A2 | lower | -1 | 2 | 1 | 5 | 5 |
| [t4] | A1A2 | A1 | lower | -1 | 2 | 1 | 5 | 5 |
| [t1] | A1A2 | A1 | upper | 1 | 2 | 1 | 5 | 5 |
| [t2] | A1A2 | A2 | upper | 1 | 2 | 1 | 5 | 5 |
| [t3] | A1A2 | A1 | upper | 1 | 2 | 1 | 5 | 5 |
| [t4] | A1A2 | A2 | upper | 1 | 2 | 1 | 5 | 5 |
| [t1] | A1 | A2 | lower | -50 | 1 | 2 | 5 | 4 |
| [t3] | A1 | A2 | lower | -100 | 1 | 2 | 5 | 4 |
| [t2] | A1 | A2 | upper | 100 | 1 | 2 | 5 | 4 |
| [t4] | A1 | A2 | upper | 100 | 1 | 2 | 5 | 4 |
| [t1] | A1 | A2 | lower | -90 | 1 | 10 | 13 | 4 |
| [t2] | A1 | A2 | upper | 900 | 1 | 10 | 13 | 4 |
| [t4] | A1 | A2 | upper | 900 | 1 | 10 | 13 | 4 |
| [t1] | A1 | A2 | lower | -99 | 1 | 100 | 103 | 4 |
| [t2] | A1 | A2 | upper | 9900 | 1 | 100 | 103 | 4 |
| [t4] | A1 | A2 | upper | 9900 | 1 | 100 | 103 | 4 |

Using the same topology and terminal distribution, but introducing a single long branch resulted in varying outcomes. While both sets exhibited equal PD values when the longest branch is located internally, the long terminal branch (terminal 1 in set A1) influenced the initial PD values, favoring the set containing that terminal (Tables 1 and 2).

Given the balanced topology and terminal distribution, swapping internal branches has no effect on PD, while swapping other branches yield the expected PD distribution: A1=0.5 / A2=0.5 (within-terminal swaps) or A1=0.33 / A1A2=0.33 / A2=0.33 (all terminals swapped) (Table 1).

Altering terminal branch lengths had a greater impact on PD than modifying internal branch lengths. In this example, the longest branch influenced the PD results, with a substantial delta (50% or greater) required in any terminal branch length to impact PD. Furthermore, the greater the relative length of the t1 branch compared to other terminals, the larger the PD difference between both sets. This suggests that results are more stable when driven primarily by a single, dominant branch length (Tables 1 and 2).

These simple examples, where the PD difference depends solely on branch length, demonstrates two distinct situations: minimal or no impact of internal branch length modifications, and impact of terminal branch length modifications. This behavior is not readily apparent from the swapping evaluation, highlighting the insufficiency of the null model in understanding the behavior of PD and its sensitivity to branch length. This emphasizes the need to evaluate the contribution of each branch length.

The R code used can be found, as a worked example, in github.com/Dmirandae/blepd/blob/primary/docs/4terminals2areas.Rmd.

### An empirical example: *Rhinoclemmys* data

In a real-world scenario, the interplay of various factors, including tree topology, the distribution of terminal taxa across sets, the specific definition of each set, and, crucially, branch lengths, might determine the PD value and its sensitivity to branch length variation.

The genus *Rhinoclemmys* genus (Testudines: Geoemydidae) comprises a group of turtles commonly known as wood turtles or river turtles, that exhibits a wide distribution, ranging from Mexico to northeastern Brazil, the “center” of diversity for this genus is located in Central America (Le and McCord, 2008). I conducted a phylogenetic diversity analysis to evaluate the effect of branch length on the chosen conservation area(s). The topology used corresponded to the Maximal likelihood total evidence analysis from Romero-Alarcon (2020) (Figure 2) and the distribution was modified from Le and McCord (2008) (Table 3).

**Figure 2.**
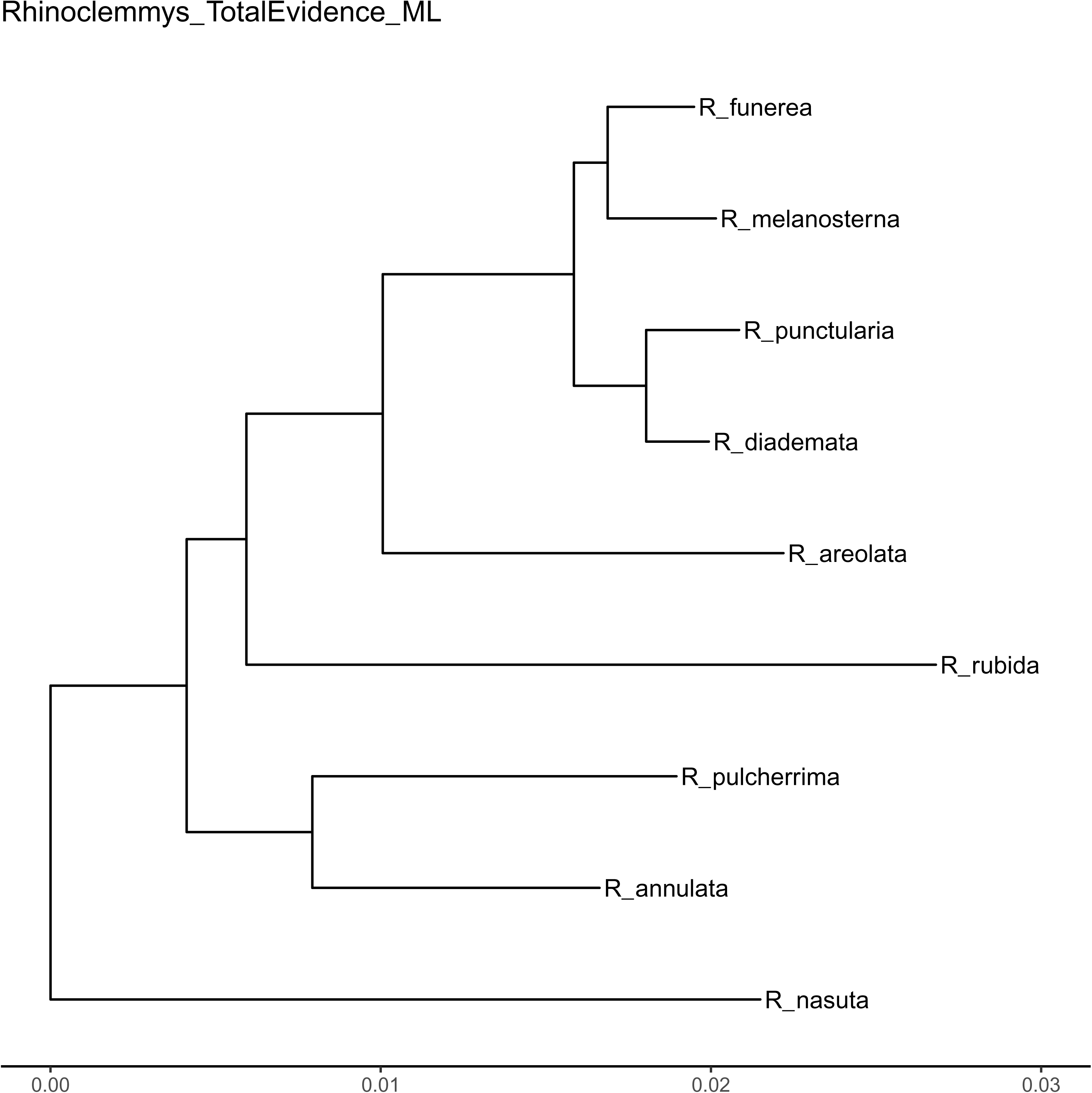
*Rhinoclemmys* phylogeny based on the total evidence maximum likelihood analysis from Alarcon-Romero (2020).

**Table 3.** *Rhynoclemmys* distribution and absolute PD values for each area. The areas of endemism follow Le and McCord (2008): Al (Atlantic Lowland), An (Amazon North), As (Amazon South), Cho (Choco), Nsa (Northern South America), Nuh (Nuclear Highland), Ph (Panamanian Herpetofauna), and Pl (Pacific Lowland). As Amazon North and Amazon South share the same species in this analysis, these two areas were combined into a single area.

| Area | annulata | areolata | diademata | funerea | melanosterna | nasuta | pulcherrima | punctularia | rubida | PD |
| --- | --- | --- | --- | --- | --- | --- | --- | --- | --- | --- |
| Al | 1 | 1 | 0 | 1 | 0 | 0 | 0 | 0 | 0 | 0.044 |
| An-As | 0 | 0 | 0 | 0 | 0 | 0 | 0 | 1 | 0 | 0.021 |
| Cho | 1 | 0 | 0 | 0 | 1 | 1 | 0 | 0 | 0 | 0.054 |
| Nsa | 0 | 0 | 1 | 0 | 0 | 0 | 0 | 0 | 0 | 0.02 |
| Nuh | 0 | 0 | 0 | 1 | 0 | 0 | 1 | 0 | 0 | 0.034 |
| Ph | 1 | 0 | 0 | 0 | 1 | 0 | 0 | 0 | 0 | 0.033 |
| Pl | 0 | 0 | 0 | 0 | 0 | 0 | 1 | 0 | 1 | 0.042 |

Cho has highest observed PD (PD=0.054, 21.86% of the total PD), followed by Al (PD=0.044, 17.82% of the total PD), which are the two richest areas (Table 3). Therefore, the main question is whether the difference in PD is a valid value for inferring the conservation status.

The branch-length histogram (Figure 3) revealed that internal branch lengths are relatively homogeneous (sd.=1.7E-3), whereas the terminal branches exhibit greater variation (sd=7.7E-3). *R. nasuta* (inhabiting Cho) and *R. rubida* (inhabiting Pl) have the longest branches (> 0.02), therefore, I expect a higher impact from swapping terminals than from swapping internal branches.

**Figure 3.**
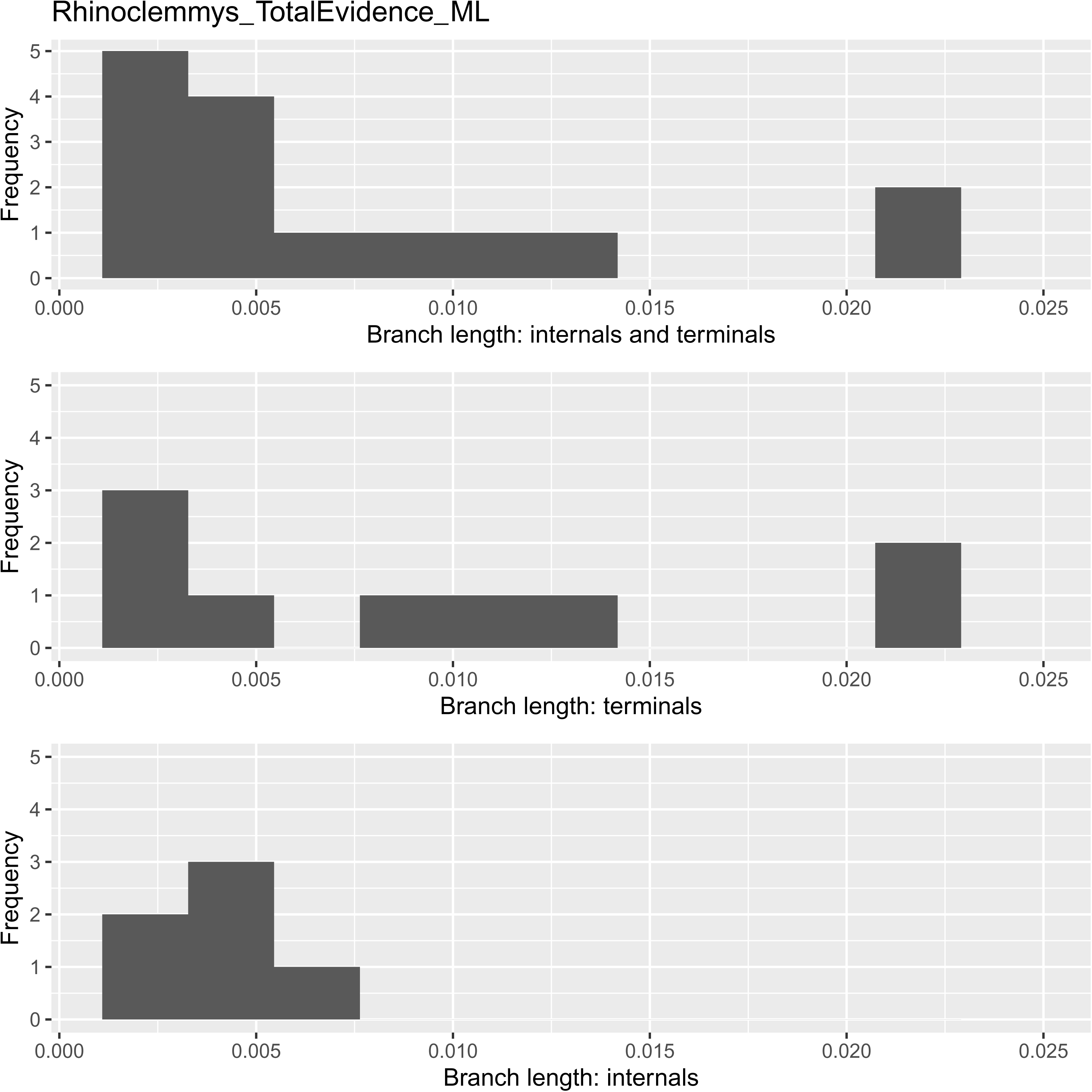
Branch length distribution for the *Rhinoclemmys* phylogeny.

Swapping the terminal or internal branch lengths using the three different argument values for the *model* shows that terminal branch lengths could be crucial for determining the selected area. While swapping internal branches has no effect, swapping terminal or all branches using “<u>allswap</u>” or “<u>uniform</u>” models suggest the inclusion of Al (Table 4).

**Table 4.** Results for SwapBL function for the *Rhinoclemmys* Example. This table shows the impact of swapping branch lengths on the selected area with the highest Phylogenetic Diversity (PD). The results are presented as the percentage of times each area was selected as optimal under different swapping scenarios: ‘terminals’: Swapping only terminal branches. ‘internals’: Swapping only internal branches. ‘all’: Swapping both terminal and internal branches. ‘allswap’: Randomly permuting all branch lengths. ‘uniform’: Resampling branch lengths from a uniform distribution.

| Branch | Model | Al | An-As | Cho | ChoNuh | Nsa | Nuh | Pl | best |
| --- | --- | --- | --- | --- | --- | --- | --- | --- | --- |
| terminals | allswap | **39.25** | 0.6 | **43.72** | 0.01 | 0.76 | 12.55 | 3.11 | Cho/Al |
| internals | allswap | 0 | 0 | **100** | 0 | 0 | 0 | 0 | Cho |
| all | allswap | **36.04** | 4.71 | **40.21** | 0 | 4.22 | 13.11 | 1.71 | Cho/Al |
| terminals | uniform | **45.64** | 0.02 | **47.96** | 0 | 0.05 | 5.86 | 0.47 | Cho/Al |
| all | uniform | **45.47** | 0.28 | **47.44** | 0 | 0.25 | 6.51 | 0.05 | Cho/Al |

Recognizing the uneven distribution of terminal branch lengths, I hypothesized that the longest branches within the Cho or Al areas might be driving these results. Therefore, to isolate the effect of branch length, I used the **evalBranch** function to evaluate the impact of modifying specific terminal branches (Table 5).

**Table 5.** Results of the evalBranch function for the *Rhinoclemmys* Example. This table shows the impact of modifying individual branch lengths on the selected area with the highest Phylogenetic Diversity (PD). node: The node identity of the branch/clade being modified. initialArea: The area with the highest PD in the initial tree. FinalArea: The area with the highest PD after the branch length modification. Approach: The type of branch length modification (“lower” for decreasing to zero, “upper” for increasing to the maximum). %Delta: The percentage change in branch length.

| node | initialArea | FinalArea | Approach | %Delta |
| --- | --- | --- | --- | --- |
| [R_pulcherrima] | Cho | Pl | upper | 116 |
| [R_diademata/R_punctularia] | Cho | An-As | upper | 1695 |
| [R_diademata] | Cho | Nsa | upper | 1710 |
| [R_punctularia] | Cho | An-As | upper | 1104 |
| [R_funerea] | Cho | Al | upper | 326 |
| [R_areolata] | Cho | Al | upper | 86 |
| [R_rubida] | Cho | Pl | upper | 59 |
| [R_nasuta] | Cho | Al | lower | -46 |

While modifying internal branches has no effect, altering some terminal branch lengths affects area selection. For example, a 46% decrease in the branch length of *R. nasuta* and an 86% increase in the branch length of *R. areolata* shifts the optimal area from Cho to Al. Similarly, a 59% increase in the branch length of *R. rubida* or a 116% increase in the branch length of *R. pulcherrima* results in a shift from Cho to Pl. Meanwhile several species such as *R. diademata*, *R. punctularia*, and the *R. diademata / R_punctularia* clade, require substantial branch length increases (over 200%) to alter the optimal area, suggesting a relatively high degree of stability in the initial area selection for these taxa.

These findings suggest that the decision to conserve only Cho is not valid and the area Al should be considered an essential component of the conservation range.

The code used to generate the analysis can be found as a worked example in github.com/Dmirandae/blepd/blob/primary/docs/RhinoclemmysExample.Rmd.

## Conclusions

The distribution of branch lengths and the impact of branch length variation cannot always be simultaneously optimized. Therefore, a compromise is necessary: the PD may not be perfectly evenly distributed, but it should be relatively insensitive to changes in branch lengths.

Since each approach (resampling or modifying individual branch lengths) evaluates a different source of information and variation, the results may sometimes appear contradictory. If branch length estimates are highly precise, resampling methods can provide reliable results. However, when there is uncertainty regarding branch length estimates, the branch length modification approach is generally more suitable. This is because branch lengths can change, and it is crucial to understand the extent to which these changes might influence the selected set.

These analyses address the critical question of whether all branch lengths contribute equally to the overall PD value or any other metric dependent on topology and branch length. Acknowledging the influence of branch length, this approach, applicable to any analysis involving branch length (including PD), will lead to a more comprehensive understanding of evolutionary history and ultimately improve decision making in evolutionary, conservation, and ecological studies.

## Acknowledgments

Laura Galvis-Tuesta tested the alpha version of the package and provided valuable insights on the development of the functions and their behavior. Viviana Romero and Iván Yesid López-Ardila provided valuable feedback on an early draft. Nicolle Saavedra proofread the manuscript. All their efforts are kindly appreciated.

